# A generative artificial intelligence approach for antibiotic optimization

**DOI:** 10.1101/2024.11.27.625757

**Authors:** Marcelo D. T. Torres, Yimeng Zeng, Fangping Wan, Natalie Maus, Jacob Gardner, Cesar de la Fuente-Nunez

## Abstract

Antimicrobial resistance (AMR) poses a critical global health threat, underscoring the urgent need for innovative antibiotic discovery strategies. While recent advances in peptide design have yielded numerous antimicrobial agents, optimizing these molecules experimentally remains challenging due to unpredictable and resource-intensive trial-and-error approaches. Here, we introduce APEX Generative Optimization (APEX_GO_), a generative artificial intelligence (AI) framework that integrates a transformer-based variational autoencoder with Bayesian optimization to design and optimize antimicrobial peptides. Unlike traditional supervised learning approaches that screen fixed databases of existing molecules, APEX_GO_ generates entirely novel peptide sequences through arbitrary modifications of template peptides, representing a paradigm shift in peptide design and antibiotic discovery. Our framework introduces a new peptide variational autoencoder with design and diversity constraints to maintain similarity to specific templates while enabling sequence innovation. This work represents the first *in vitro* and *in vivo* experimental validation of generative Bayesian optimization in any setting. Using ten de-extinct peptides as templates, APEX_GO_ generated optimized derivatives with enhanced antimicrobial properties. We synthesized 100 of these optimized peptides and conducted comprehensive *in vitro* characterizations, including assessments of antimicrobial activity, mechanism of action, secondary structure, and cytotoxicity. Notably, APEX_GO_ achieved an outstanding 85% ground-truth experimental hit rate and a 72% success rate in enhancing antimicrobial activity against clinically relevant Gram-negative pathogens, outperforming previously reported methods for antibiotic discovery and optimization. In preclinical mouse models of *Acinetobacter baumannii* infection, several AI-optimized molecules—most notably derivatives of mammuthusin-3 and mylodonin-2—exhibited potent anti-infective activity comparable to or exceeding that of polymyxin B, a widely used last-resort antibiotic. These findings highlight the potential of APEX_GO_ as a novel generative AI approach for peptide design and antibiotic optimization, offering a powerful tool to accelerate antibiotic discovery and address the escalating challenge of AMR.

## Introduction

Antimicrobial resistance (AMR) poses a significant and escalating global health threat, necessitating the discovery of new antibiotics to combat resistant pathogens^1^. Peptide-based therapeutics, particularly antimicrobial peptides (AMPs), have emerged as promising candidates due to their broad-spectrum activity and unique mechanisms of action^1–3^. However, designing and optimizing such peptides is a complex task^4^, often hindered by the vastness of the peptide sequence space and the unpredictability of experimental trial-and-error methods.

Recent advances in molecular de-extinction^5,6^ have opened new avenues for antibiotic discovery by identifying and characterizing biological molecules from the past. This approach has uncovered a new sequence space that may harbor potent antimicrobial agents capable of addressing modern challenges like AMR. Leveraging this concept, we previously developed APEX^6^, a deep learning-based functional predictor of antimicrobial activity, to mine all extinct organisms known to science, leading to the discovery of numerous new antibiotics.

Despite these successes, optimizing the antimicrobial properties of peptides remains challenging. Traditional methods rely heavily on iterative experimental modifications, which are time-consuming and resource intensive^7–9^. Thus, there is a pressing need for efficient optimization approaches that can navigate the vast peptide sequence space and enhance desirable properties such as potency, spectrum of activity, and safety profiles.

In this study, we introduce APEX Generative Optimization (APEX_GO_), a generative artificial intelligence framework that integrates deep generative modeling with Bayesian optimization (BO) to streamline peptide design and optimization. APEX_GO_ employs a transformer-based variational autoencoder (VAE) to map peptide sequences into a continuous latent space, transforming the discrete optimization problem into a tractable continuous one. Here, we trained a new generative VAE model for peptide sequences. By incorporating BO techniques, including our own methods like LOL-BO^10^, APEX_GO_ efficiently explores this latent space to identify novel peptides with enhanced antimicrobial properties. Combining generative AI and Bayesian optimization represents a significant methodological departure from recent successful prior work that uses supervised deep learning to virtually screen large but fixed databases of molecules^11–13^. APEX_GO_ instead searches over arbitrary peptide modifications, and can suggest peptides that would not exist in any known database. Different from prior work^10,14^ in generative Bayesian optimization, we seek to design optimized derivatives of existing templates, suspecting that de-extinct peptides are more likely to evade antibiotic resistance. We augment latent space BO with design and diversity constraints, and a per-template fine-tuned peptide VAE. Notably, our work represents the first ground-truth *in vitro* or *in vivo* experimental validation of generative Bayesian optimization.

Using ten de-extinct peptides as templates, we applied APEX_GO_ to generate optimized derivatives with improved antimicrobial activity (**Supplementary Table 1**). The templates were selected based on their varied activity profiles and potential for enhancement. We synthesized 100 optimized peptides and conducted comprehensive *in vitro* characterizations, including assessments of antimicrobial efficacy against clinically relevant pathogens, mechanism of action studies, secondary structure analyses, and cytotoxicity evaluations.

Notably, the optimized peptides demonstrated significant improvements in antimicrobial potency, including activity against multidrug-resistant strains. In preclinical mouse models of *Acinetobacter baumannii* infection, several optimized molecules—specifically those derived from mammuthusin-3 and mylodonin-2—exhibited potent anti-infective activity comparable to or exceeding that of polymyxin B, a last-resort antibiotic. These findings highlight the effectiveness of APEX_GO_ as a novel generative AI approach for antibiotic optimization, opening new avenues to address AMR.

## Results and Discussion

### APEX_GO_: integrating deep learning with Bayesian optimization

We recently developed APEX, a deep learning approach to predict antibiotic function from amino acid sequence^6^. APEX efficiently predicts the minimum inhibitory concentrations (MICs) of peptides against a variety of Gram-negative and Gram-positive bacterial pathogens (see **APEX** in **Methods** section). While APEX successfully discovered antibiotic molecules from extinct organisms, its discriminative learning nature limits its capacity for *de novo* design or property optimization. Moreover, the vastness of the peptide sequence space renders exhaustive virtual screening infeasible.

To address these challenges, we developed APEX_GO_, which integrates APEX with black-box Bayesian optimization (BO) and generative modeling methods to perform constrained template-based optimization of antibiotic activity in peptide sequences (**Fig. 1a**). APEX_GO_ employs a transformer-based variational autoencoder (VAE)^15,16^ to map peptide sequences into a continuous latent space, effectively transforming the discrete optimization problem into a tractable continuous one. This approach enables efficient exploration of the sequence space to identify molecules with improved antimicrobial properties.

**Figure 1.**
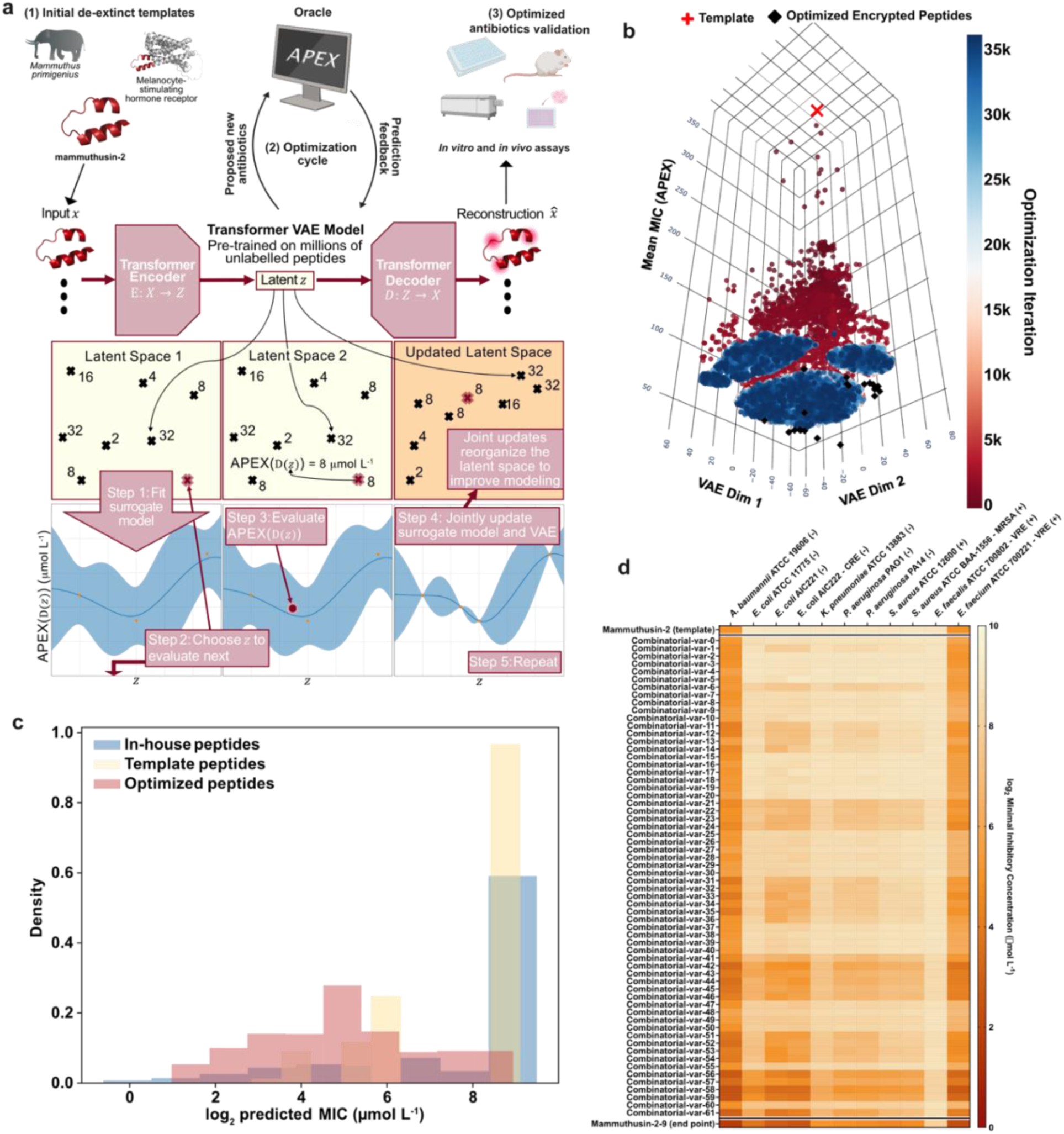
APEX_GO_: computational workflow and results. **(a)** The overall workflow of APEX_GO_ demonstrates de-extinction, optimization and validation. The workflow shows how the peptide VAE is jointly updated with the surrogate model during optimization. **(b)** Visualization of latent space optimization in 2D using t-SNE. The red cross indicates the template peptide sequence with a high APEX-predicted mean MIC. Black diamonds represent the optimized peptides identified by APEX_GO_. Scattered points correspond to latent space samples evaluated during optimization, with the color gradient indicating the progression of optimization iterations from early (red) to later stages (blue). **(c)** Comparison of log_2_ transformed MIC predictions by APEX for three peptide groups: the in-house peptides used to train APEX, the ten peptide templates selected for optimization, and the optimized peptides generated by APEX_GO_. Box plots display the distribution of predicted MIC values, demonstrating a significant shift toward lower MICs in the optimized peptides, indicative of enhanced predicted antimicrobial activity. **(d)** Heatmap showing the predicted MICs for a specific template peptide (top row), the final optimized peptide (bottom row), and intermediate peptide variants (middle rows) obtained by systematically reverting individual amino acid substitutions back to the template sequence. This analysis illustrates that the amino acid changes introduced by APEX_GO_ are computationally necessary for improving antimicrobial potential.

In APEX_GO_, the generation process involves sampling latent space points and decoding them into peptide sequences using the VAE decoder. The evaluation process uses APEX to predict the antimicrobial potential of the generated peptides. A surrogate model, implemented as a parametric Gaussian process regressor (PPGPR)^17^, models the correlation between the latent space points and the APEX-predicted MICs. The BO algorithm utilizes this surrogate model to propose new latent space points likely to decode into peptides with enhanced antimicrobial activity. This iterative process continues until peptides with optimized properties are identified.

Key features of APEX_GO_ include:

1. **Bayesian Optimization over Adaptive Latent Spaces**: By training a VAE that maps peptides to a continuous latent space, we convert a discrete optimization problem into a continuous one, facilitating optimization. Periodic joint updates of the VAE and surrogate model encourage the latent space to reorganize, clustering peptides with similar MICs closer together, thereby enhancing modeling and optimization.
2. **Multiple Trust Regions**: To prevent over-exploration in the high-dimensional latent space, we implement trust region Bayesian optimization^18^. This method defines hyper-rectangular regions centered on the best points observed so far, adapting their size based on optimization progress. By using multiple trust regions, we simultaneously optimize multiple distinct peptides in a single run.

### Template-based antimicrobial peptide design and optimization

In this work, we focused on template-based AMP optimization, using ten peptides mined from the proteomes of extinct organisms as templates. These templates were selected based on their varied activity profiles, ranging from selective to broad-spectrum effects. Importantly, none of the template de-extinct peptides were highly active (MIC ≤ 16 μmol L⁻¹), allowing room for optimization.

APEX_GO_ performed constrained peptide sequence optimization, ensuring that each optimized sequence maintained at least 75% sequence similarity to its template (see **Methods** section). The optimization aimed to improve antimicrobial activities against either seven Gram-negative pathogen strains or all eleven strains (including Gram-positive bacteria) that APEX predicts (see **APEX** in **Methods** section).

Starting from the selected template sequences with relatively high average MICs against Gram-negative bacteria, APEX_GO_ generated sequences with progressively lower predicted MICs as optimization progressed (**Fig. 1b**). Compared to the MIC distributions of the APEX training peptides and the template peptides, the optimized peptides showed a significant shift toward lower experimental MICs, indicating enhanced predicted antimicrobial activities (**Fig. 1c**). Statistical analysis confirmed that the prediction improvements were significant (p-values of one-sided Mann-Whitney U test were 1.19e-309 and 1.12e-35 when comparing optimized peptides to inhouse and template peptides, respectively).

To assess the contribution of each amino acid change proposed by APEX_GO_, we generated intermediate variants by systematically reverting individual mutations in the optimized sequences back to the template amino acids (**Fig. 1d**). By evaluating these variants with APEX and ranking them based on predicted MICs, we found that the optimized peptides were almost always ranked first (mean ranking of 1.23; standard deviation of 0.42), suggesting that nearly all amino acid changes proposed by APEX_GO_ were essential for improving antimicrobial potential.

### Ground-truth in vitro antimicrobial activity of optimized peptides

To validate the predictive accuracy of APEX_GO_, we synthesized and tested two sets of optimized peptides per de-extinct template: (i) five peptides predicted to be active against Gram-negative bacteria and (ii) five peptides predicted to be broad-spectrum. A total of 100 optimized peptides were synthesized and experimentally tested for antimicrobial activity against 11 clinically relevant bacterial pathogens, including strains resistant to conventional antibiotics (**Fig. 2**).

**Figure 2.**
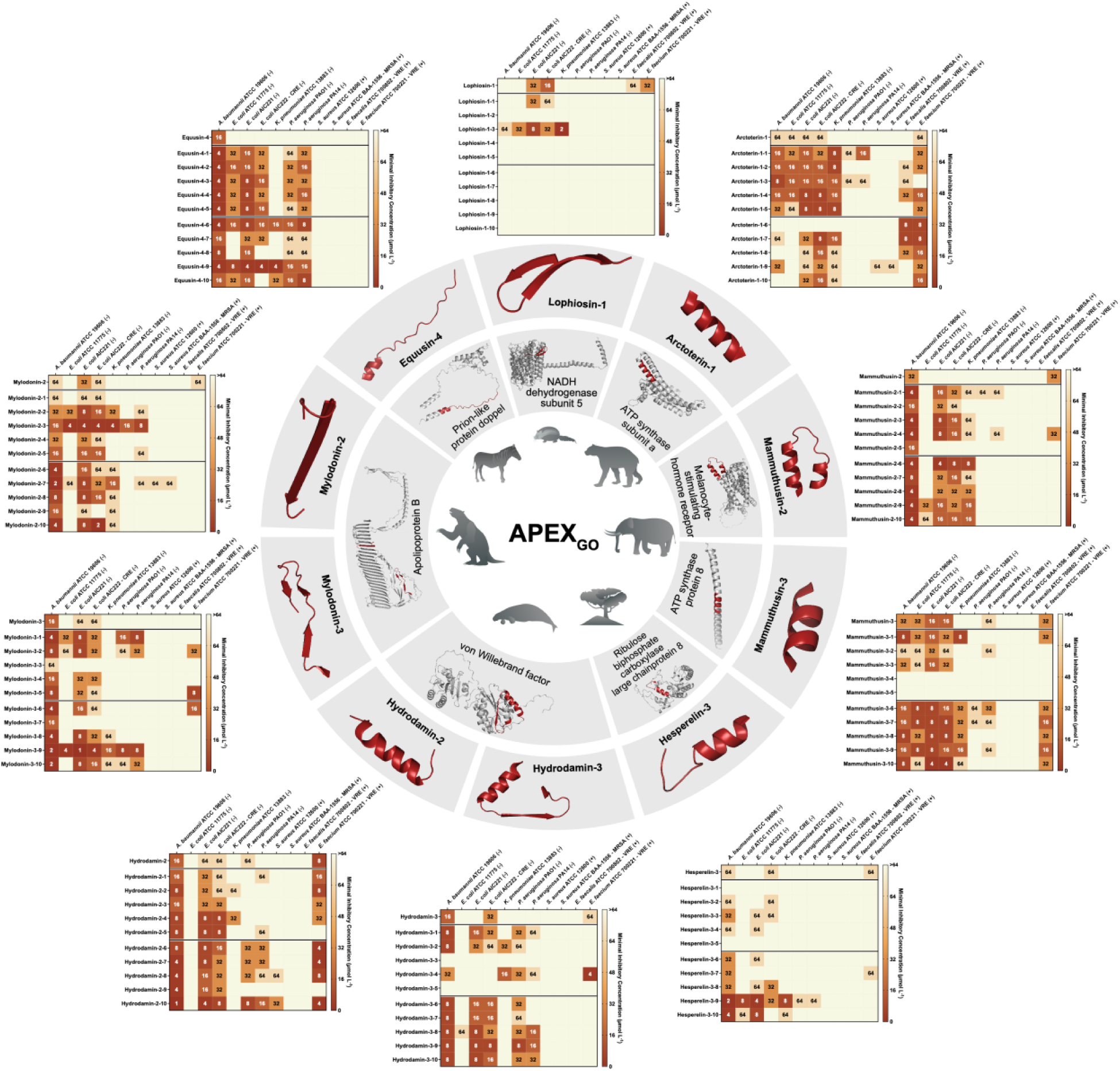
Antimicrobial activity of peptides optimized by APEX_GO_. Heat map displaying the minimum inhibitory concentrations (MICs, in μmol L⁻¹) of peptides optimized by APEX_GO_ against clinically relevant bacterial pathogens, including four strains resistant to conventional antibiotics. The MICs were determined using the broth microdilution method, and the heat map presents the mode MIC values from three independent biological replicates for each condition. Lower MIC values (indicated by darker shades) represent higher antimicrobial potency. The optimized peptides demonstrate significant improvements in antimicrobial efficacy compared to their parent templates, validating the effectiveness of APEX_GO_ in generating peptides with enhanced activity against both standard and multidrug-resistant bacterial strains.

Of the 100 synthesized peptides, 86 exhibited significant antimicrobial activity (MIC ≤ 64 μmol L^-1^) against at least one bacterial strain, resulting in an 86% hit rate —greatly surpassing the 59% hit rate achieved by APEX alone^6^. Pearson and Spearman correlation coefficients between experimentally determined MICs and APEX predictions for the 100 optimized sequences were 0.463 and 0.462, respectively, underscoring APEX’s significant predictive power and the effectiveness of using APEX as the oracle for BO.

Comparing the mean experimentally determined MICs between optimized peptides and their corresponding templates, we observed that 68% of the optimized peptides showed improved antimicrobial activity after optimization with APEX_GO_ (**Fig. 2**). When considering the Gram-negative strains only, the improvement rate reached 72%. Template-wise and bacterial strain-wise analyses (i.e., in terms of average mean MIC improvement rate and average MIC log-fold change) indicated that APEX_GO_ was particularly effective at optimizing peptides against Gram-negative pathogens (85% hit rate), consistent with APEX’s stronger predictive performance for these bacteria^6^. That being said, peptides optimized for broad-spectrum activity had significantly lower MIC distributions than those optimized specifically for Gram-negative bacteria (p-value of one-sided Mann-Whitney U test: 0.006), highlighting the value of broad-spectrum optimization in APEX_GO_.

Specific observations include modifications to equusin-4 derivatives, such as replacing histidine with isoleucine at position 18 (H18I), which improved activity against Gram-negative strains, particularly *Acinetobacter baumannii*, *Escherichia coli*, and *Pseudomonas aeruginosa*. Arctoterin-1 optimized peptides showed increased activity against Gram-negative bacteria, with substitutions such as histidine to glycine at position 6 (H6G) enhancing activity against *S. aureus*. Positively charged modifications in lophiosin-1 at the N-terminus maintained antimicrobial activity, while negatively charged substitutions failed to improve broad-spectrum efficacy. In mylodonin-2, substitutions like threonine to arginine or tryptophan at position 10 (T10R or T10W) enhanced activity against *Klebsiella pneumoniae*, especially when combined with amphiphilic N-terminal modifications. For mammuthusins 2 and 3 peptides, increases in normalized hydrophobic moment generally correlated with improved activity, except in specific cases where crucial residues were altered, such as isoleucine at position 6 in mammuthusin-3-5. Hydrodamin-2 peptides showed increased or maintained activity with various modifications, except for hydrodamin-2-10, which had a unique tyrosine to tryptophan substitution at position 9 (Y9W). Finally, hesperelin-3 peptides lost activity with the introduction of a proline residue in place of glycine at position 6 (G6P), whereas the most effective broad-spectrum peptides preserved the arginine at position 1 (R1).

### Secondary structure of optimized peptides

Since the optimized peptides sometimes differed significantly from the templates in terms of physicochemical descriptors and antimicrobial activity profiles, we investigated whether the optimization led to changes in their secondary structure compared to the templates. To assess the secondary structure of the peptides, we exposed them to three different media (**Fig. 2a-j, Supplementary Figs. 1-3**): water, helix-inducing medium^19^ (trifluoroethanol in water, 3:2, v:v), and membrane-mimicking environment (Sodium dodecyl sulfate, SDS, at 10 mmol L^-1^).

Equusin-4 derivatives exhibiting higher helicity (**Fig. 3a, Supplementary Figs. 1a-c, 3**), particularly in the helix-inducing medium, were more active, especially against *K. pneumoniae* (**Fig. 2**). In contrast, arctoterin-1 derivatives (**Fig. 3b, Supplementary Figs. 1d-f, 3**) showed no clear correlation between helicity and activity; the optimized peptides that were more active against Gram-negative bacteria were less helical. Lophiosin-1 (**Fig. 3c, Supplementary Figs. 1g-i, 3**) and mylodonin-2 (**Fig. 3e, Supplementary Figs. 1m-o, 3**) derivatives were mostly β-structured or unstructured. In these cases, lower helicity correlated with higher activity, whereas more α-helical derivatives were inactive. Although mylodonin-3 derivatives were primarily unstructured, peptides with slightly higher β-sheet content were more active (**Fig. 3d, Supplementary Figs. 1j-l, 3**).

**Figure 3.**
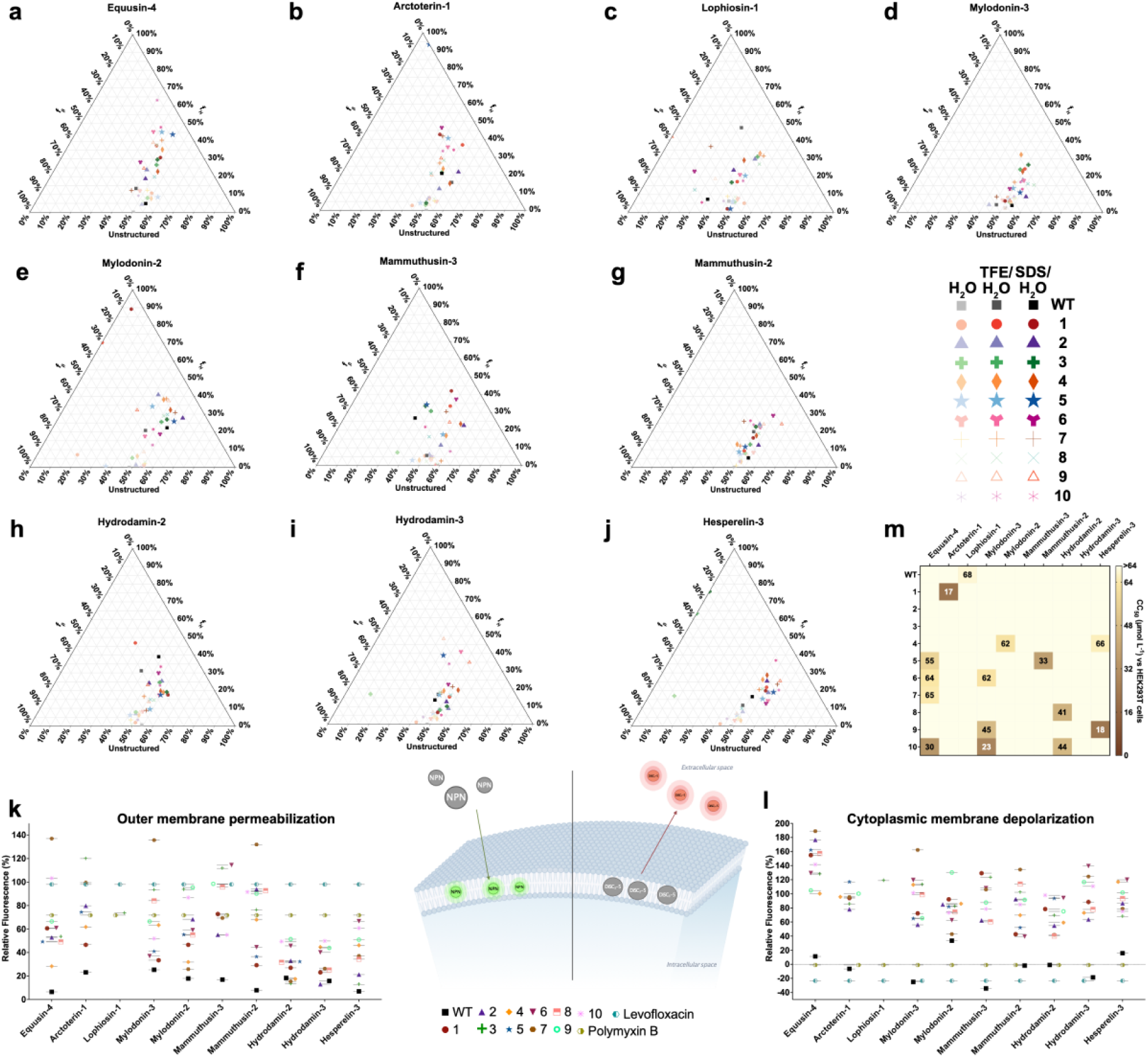
Secondary structure, cytotoxicity, and mechanism of action of templates and peptides optimized by APEX_GO_. **(a-j)** Ternary plots showing the percentage of secondary structure for each peptide (at 50 μmol L^-1^) in three different solvents: water, 60% trifluoroethanol (TFE) in water, and Sodium dodecyl sulfate (SDS, 10 mmol L^-1^) in water. Secondary structure fractions were calculated using the BeStSel server^28^. **(k)** Outer membrane permeabilization of *Acinetobacter baumannii* ATCC 19606 induced by the peptides, assessed using the fluorescent probe 1-(N-phenylamino)naphthalene (NPN). The values represent the end-point relative fluorescence units (RFU) obtained after nonlinear fitting, compared to the baseline of the untreated control (buffer + bacteria + NPN), and benchmarked against the template peptides and the antibiotics polymyxin B and levofloxacin. **(l)** Cytoplasmic membrane depolarization of *A. baumannii* ATCC 19606 induced by the peptides, evaluated using the potentiometric fluorescent probe 3,3′-dipropylthiadicarbocyanine iodide (DiSC₃-5). The values displayed represent the end-point RFU obtained after nonlinear fitting, compared to the baseline of the untreated control (buffer + bacteria + DiSC₃-5), and benchmarked against the template peptides and the antibiotics polymyxin B and levofloxacin. **(m)** Cytotoxic concentrations leading to 50% cell lysis (CC₅₀) in human embryonic kidney (HEK293T) cells, determined by interpolating dose–response data using a nonlinear regression curve. All experiments were performed in three independent replicates.

Mammuthusin-2 (**Fig. 3g, Supplementary Figs. 2d-f, 3**) and mammuthusin-3 (**Fig. 3f, Supplementary Figs. 2a-c, 3**) derivatives exhibited low helicity with a slight increase in α-helical structure in lipid bilayers; however, no clear correlation with antimicrobial activity was observed. Similarly, hydrodamin-2 (**Fig. 3h, Supplementary Figs. 2g-i, 3**) and hydrodamin-3 (**Fig. 3i, Supplementary Figs. 2j-l, 3**) derivatives showed a slight increase in α-helicity in membrane-mimicking environments, but their antimicrobial activity did not correlate with secondary structure. Hesperelin-3 derivatives were mostly unstructured (**Fig. 3j, Supplementary Figs. 2m-o, 3**). An exception was one derivative that showed a significant increase in α-helicity in the presence of lipid bilayers; however, this did not strongly correlate with activity. Overall, no consistent trend was observed between secondary structure and antimicrobial activity across the peptides studied.

### Mechanism of action of optimized peptides

The bacterial membrane is a common target for most AMPs, where they engage in non-specific interactions with the lipid bilayer^8^. The antimicrobial activity of AMPs is influenced by their amino acid composition, distribution, and various physicochemical characteristics such as amphiphilicity and hydrophobicity. To investigate the underlying mechanisms by which the AI-optimized peptides predicted by APEX_GO_ kill bacteria, we tested whether differences in the composition of the derivatives —resulting from substitutions and/or insertions made to the original sequences—would affect their mechanism of action. For these assays, we selected *A. baumannii* ATCC 19606, which was the most sensitive strain in our MIC assays.

First, we tested whether the peptides permeabilized the bacterial outer membrane using 1-(N-phenylamino)naphthalene (NPN) assays (**Fig. 3k and Supplementary Fig. 4**). NPN, a lipophilic dye, fluoresces faintly in aqueous solutions but fluoresces significantly more when it encounters lipidic environments such as bacterial membranes. NPN is a lipophilic dye that fluoresces faintly in aqueous solutions but exhibits significantly increased fluorescence in lipidic environments such as bacterial membranes. NPN can penetrate the bacterial outer membrane only if it is disrupted or compromised. Among the mylodonin-3 derivatives, mylodonin-3-7 was the only analog that exhibited superior permeabilization compared to both control antibiotics (**Fig. 3k and Supplementary Fig. 4g,h**). Mammuthusin-3 derivatives -3, -8, and -9 also showed slightly better permeabilization than the controls (**Fig. 3k and Supplementary Fig. 4k,l**), although not as effective as mylodonin-3-7. Within the mammuthusin-2 family, mammuthusin-2-7 had the highest permeabilization activity, with derivatives -2, -8, -9, and -10 being slightly more effective than the antibiotics (**Fig. 3k and Supplementary Fig. 4m,n**). Arctoterin-1-9 was the most effective permeabilizer among its family, while derivatives -3 and -7 demonstrated slight permeabilization improvements over the controls. (**Fig. 3k and Supplementary Fig. 4c,d**) Equusin-4-7 was the strongest permeabilizer among its derivatives, with equusin-4-10 showing marginally better depolarization than the antibiotics (**Fig. 3k and Supplementary Fig. 4a,b**). In contrast, the only lophiosin-3 derivative active against *A. baumannii* was not an effective permeabilizer (**Fig. 3k and Supplementary Fig. 4e,f**). Additionally, none of the mylodonin-2 (**Fig. 3k and Supplementary Fig. 4i,j**), hesperelin-3 (**Fig. 3k and Supplementary Fig. 4s,t**), hydrodamin-3 (**Fig. 3k and Supplementary Fig. 4q,r**), or hydrodamin-2 (**Fig. 3k and Supplementary Fig. 4o,p**) derivatives demonstrated notable permeabilization activity.

Next, we tested whether the optimized peptides depolarized the cytoplasmic membrane of *A. baumannii* (**Fig. 3l and Supplementary Fig. 5**). We used the potentiometric fluorophore 3,3′-dipropylthiadicarbocyanine iodide (DiSC_3_-5), whose fluorescence is suppressed by its accumulation and aggregation within the cytoplasmic membrane. Upon disturbances in the transmembrane potential of the cytoplasmic membrane, this fluorophore migrates to the outer environment and emits fluorescence. Polymyxin B was used as a positive control in these experiments, as it is a depolarizer that also permeabilizes and damages bacterial membranes. All peptides tested demonstrated stronger depolarization activity compared to the antibiotics polymyxin B and levofloxacin. Among the mylodonin-3 derivatives, mylodonin-3-7 was the most effective depolarizer of *A. baumannii*, followed closely by mylodonin-3-3, -4, -6, -8, and -10 (**Fig. 3l and Supplementary Fig. 5g,h**). Mylodonin-2-9 was the top depolarizer among the mylodonin-2 analogs (**Fig. 3l and Supplementary Fig. 5i,j**). Mammuthusin-3 derivatives -1, -3, -6, and -7 were slightly better depolarizers than the others (**Fig. 3l and Supplementary Fig. 5k,l**), while mammuthusin-2-7 exhibited the highest depolarization activity within its family, followed by derivatives -2, -3, -8, -9, and -10 (**Fig. 3l and Supplementary Fig. 5m,n**). Lophiosin-1-3 (**Fig. 3l and Supplementary Fig. 5e,f**) and several hydrodamin-3 analogs (-6, -7, -9, and -10) were also effective depolarizers (**Fig. 3l and Supplementary Fig. 5q,r**). Among the hydrodamin-2 derivatives, -5, -6, and -10 showed the best activity (**Fig. 3l and Supplementary Fig. 5o,p**), although slightly less than the top performers from other groups. Hepserelin-3 derivatives -4 and -6, and arctoterin-1-5 were the strongest depolarizers in their respective families (**Fig. 3l and Supplementary Fig. 5s,t**). Equusin-4 derivatives, particularly -2 and -7, exhibited uniformly high depolarization activity (**Fig. 3l and Supplementary Fig. 5a,b**). Overall, the best depolarizers from each group displayed comparable efficacy, with hydrodamin-2 derivatives being slightly less potent than the rest.

### Cytotoxicity assays

All derivatives were tested for cytotoxic activity against human embryonic kidney (HEK293T) cells and compared to their templates (**Fig. 3m**). This assay is widely used to assess the toxicity of antimicrobials because the results are highly reproducible^20–22^. Of the peptides tested, 84 displayed no significant cytotoxicity at the concentration range tested (4-64 μmol L^-1^). Among the equusin-4 derivatives, only 5, 6, 7, and 10 showed mild to moderate toxicity, while the most active derivatives were non-toxic. Arctoterin-1-1, the most active and broad-spectrum derivative, was the only cytotoxic peptide in its family. Interestingly, while the lophiosin-1 template was cytotoxic, all its derivatives were non-toxic. Most mylodonin-2 derivatives were non-toxic except for mylodonin-2-4 at high concentrations.

Mylodonin-3 derivatives optimized for broad-spectrum activity exhibited higher toxicity compared to those optimized for Gram-negatives, with mylodonin-3-9 and 10 being the most toxic, though still at concentrations 10 to 20 times higher than their MICs. Among the mammuthusin-2 derivatives, only mammuthusin-2-5, which had lower antimicrobial activity, was toxic. Mammuthusin-3 and hydrodamin-3 derivatives were non-toxic within the tested concentration ranges. Hydrodamin-2 derivatives with double or triple tryptophan substitutions, optimized for broad-spectrum activity, showed higher toxicity than those targeting Gram-negatives. Hesperelin-3 derivatives were generally non-toxic, except for hesperelin-3-9, which had a MIC 5 to 10 times lower than its cytotoxic concentration. Overall, the cytotoxicity varied across peptide families and was often related to the specific optimizations made for antimicrobial activity.

### Anti-infective efficacy of optimized peptides in animal models

To evaluate the *in vivo* anti-infective efficacy of the most active optimized derivatives from five highly active template peptides, we used two preclinical mouse models: skin abscess^22–25^ and intramuscular thigh infection^25–27^. Five de-extinct compounds active against *A. baumannii* and their respective most active, non-toxic AI-optimized derivatives were tested with a single dose at their MIC concentration after the infection was established. The peptides had a wide range of MIC values (4-64 μmol L^-1^) when tested *in vitro* against *A. baumannii*: equusin-4 and equusin-4-6 (MIC values of 16 and 4 μmol L^-1^, respectively) from the extinct zebra *Equus quagga boehmi*; arctoterin-1 and arctoterin-1-3 (MIC values of 64 and 8 μmol L^-1^, respectively) from the extinct bear *Arctotherium sp.*; mammuthusin-3 and mammuthusin-3-6 (MIC values of 32 and 16 μmol L^-1^, respectively) from the woolly mammoth *Mammuthus primigenius*; hydrodamin-3 and hydrodamin-3-9 (MIC values of 16 and 8 μmol L^-1^, respectively) from the extinct manatee *Hydrodamalis gigas*; and mylodonin-2 and mylodonin-2-3 (MIC values of 64 and 16 μmol L^-1^, respectively) from the extinct giant sloth *Mylodon darwinii*.

In the skin abscess infection model, mice were infected with a bacterial load of 5×10^5^ cells of *A. baumannii* (**Fig. 4a**). Each peptide was administered as a single dose over the infected area. After two days, bacterial counts revealed that all tested peptides, except arctoterin-1, arctoterin-1-3, and hydrodamin-3-9, significantly reduced the bacterial load by up to 4 orders of magnitude. In most cases, the optimized derivatives demonstrated comparable activity to their template peptides, with the exception of mylodonin-2-3, which showed greater activity than its parent peptide. These findings underscore the potent anti-infective properties of these molecules against *A. baumannii* infections and the efficacy of our APEX_GO_ optimization model in generating high-activity peptides. After four days, the optimized derivatives exhibited slightly higher activity than their parent compounds, with equusin-4-6 and mylodonin-2-3 showing antibacterial efficacy similar to that of widely used antibiotics polymyxin B and levofloxacin (**Fig. 4b**). No variations in weight, damage to the skin tissue, or other harmful consequences induced by the molecules or their derivatives were observed in the mice throughout our experiments (**Supplementary Fig. 6a**).

**Figure 4.**
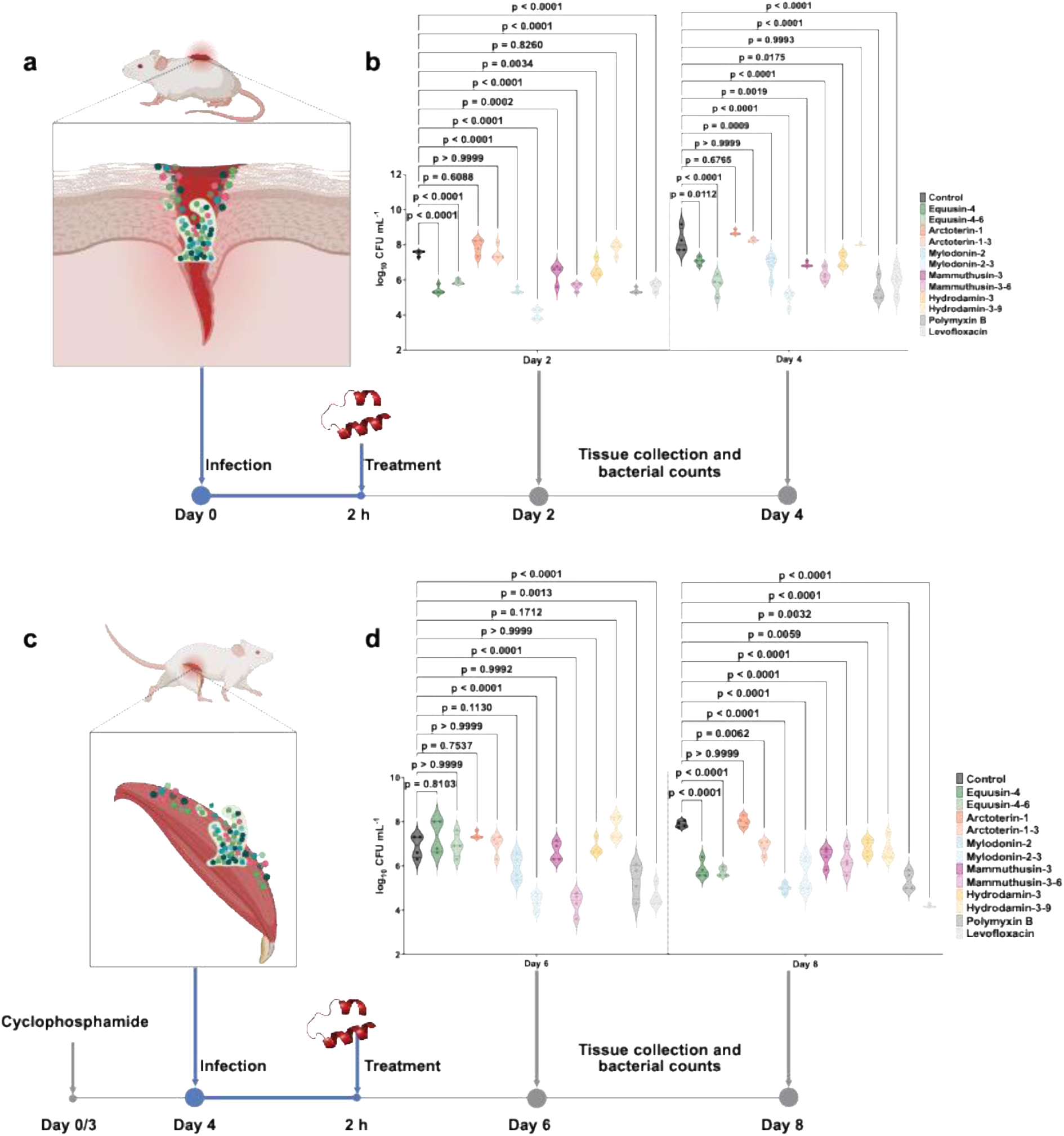
Anti-infective activity of templates and APEX_GO_-optimized peptides in animal models. **(a)** Schematic representation of the skin abscess mouse model used to assess the anti-infective activity of the designed peptides against *A. baumannii* ATCC 19606. **(b)** Comparison of the antibacterial efficacy between five parent de-extinct peptides (equusin-4, arctoterin-1, mylodonin-2, mammuthusin-3, and hydrodamin-3) and their most active, non-toxic APEX_GO_-optimized derivatives (equusin-4-6, arctoterin-1-3, mylodonin-2-3, mammuthusin-3-6, and hydrodamin-3-9, respectively). Peptides were administered as a single dose at their respective MICs post-infection (n = 4 per group). Two days after infection, several optimized peptides— including equusin-4-6, mylodonin-2-3, and mammuthusin-3-6—significantly reduced bacterial loads, demonstrating activity comparable to the control antibiotics polymyxin B and levofloxacin. Notably, mylodonin-2-3 was the most effective, reducing the infection by four orders of magnitude compared to the untreated control and outperforming the antibiotics by one order of magnitude. After four days, the optimized peptides generally exhibited higher anti-infective activity than their parent peptides, with equusin-4-6 and mylodonin-2-3 maintaining efficacy equivalent to the control antibiotics. **(c)** Schematic representation of the deep thigh infection mouse model, where neutropenic mice were infected with *A. baumannii* ATCC 19606 and treated intraperitoneally with a single dose of the same parent peptides and their APEX_GO_-optimized derivatives (n = 4 per group). **(d)** Two days post-infection and treatment (day 6), optimized peptides such as mylodonin-2-3 and mammuthusin-3-6 demonstrated significant antibacterial activity, reducing bacterial counts by three orders of magnitude compared to the untreated control group and matching the efficacy of the positive control antibiotics. Four days after treatment (day 8), all optimized peptides continued to show anti-infective activity, except for the parent peptide arctoterin-1. Statistical significance in panels **(b)** and **(d)** was determined using one-way ANOVA followed by Dunnett’s test; p-values are indicated in the graphs. In the violin plots, the center line represents the mean, the box limits indicate the first and third quartiles, and the whiskers (minima and maxima) extend to 1.5 × the interquartile range. Panel illustrations in **(a)** and **(c)** were created with BioRender.com.

In the murine deep thigh infection model, the efficacy of the same optimized derivatives and their templates was assessed following the establishment of an intramuscular infection (**Fig. 4c**). This well-established preclinical model is suited for evaluating the translatability of potential antibiotics. Mice were rendered neutropenic with cyclophosphamide before intramuscular injection of 9×10^5^ *A. baumannii* cells (**Fig. 4d**). A single dose of each compound at its MIC was administered intraperitoneally. Two days post-treatment, only mylodonin-2-3 and mammuthusin-3-6 demonstrated activity comparable to the positive control antibiotics, polymyxin B and levofloxacin, reducing bacterial counts by 3 orders of magnitude. Four-days post-treatment, most peptides, except for arctoterin-1, showed bacteriostatic activity reducing the bacterial load by 2 orders of magnitude compared to the untreated control group (**Fig. 4d**). No toxicity was observed in treated mice, as indicated by stable weight monitoring (**Supplementary Fig. 6b**).

These robust *in vivo* results demonstrate that the optimized derivatives possess potent anti-infective efficacy under physiologically relevant conditions. The optimized peptides not only improved upon their template counterparts but also performed comparably to widely used antibiotics, highlighting their potential as effective antimicrobial agents.

## Conclusion

This study demonstrates the power of integrating generative artificial intelligence approaches for antibiotic optimization. We present APEX_GO_, which successfully enhanced the activity of peptides mined from extinct organisms and identified derivatives with potent antimicrobial efficacy against high-priority bacterial pathogens, including multidrug-resistant strains. The optimized compounds exhibited favorable *in vitro* and *in vivo* profiles, with some outperforming conventional antibiotics in preclinical models.

Our findings underscore the potential of combining deep learning and Bayesian optimization to accelerate antibiotic discovery. APEX_GO_ represents a significant advancement in peptide design, streamlining the optimization process and opening promising avenues for developing new antibiotics and other therapeutics. Translating antimicrobial agents to clinical use involves optimizing properties beyond antimicrobial activity, such as stability, bioavailability, and safety. In future work, we will focus on extending APEX_GO_ to include multi-property optimization.

## Methods

### APEX 1.1

APEX^6^ is a deep learning-based model that takes peptide sequences as inputs and predicts minimum inhibitory concentration (MIC, in μmol L^-1^) against *A. baumannii* ATCC 19606, *E. coli* ATCC 11775, *E. coli* AIC221, *E. coli* AIC222, *K. pneumoniae* ATCC 13883, *P. aeruginosa* PAO1, *P. aeruginosa* PA14, *S. aureus* ATCC 12600, methicillin-resistant *S. aureus* ATCC BAA-1556, vancomycin-resistant *E. faecalis* ATCC 700802, and vancomycin-resistant *E. faecium* ATCC 700221. APEX 1.1^29^ used a hybrid of recurrent and attention neural networks to extract peptide level features that are useful for antimicrobial prediction from physicochemical and biomedical properties of amino acid sequences and was jointly trained by publicly available antimicrobial peptides^30–32^ and our internal peptide antimicrobial activity data. Through wet-lab experimental validation, we demonstrated the reliability of APEX 1.1 on finding antimicrobial peptides. In the APEX_GO_ framework, APEX is served as the oracle function and provides MICs of peptides to guide the Bayesian optimization to propose new sequences with improved antimicrobial activities.

### Black-box and Bayesian optimization

In black-box optimization, we aim to optimize an *oracle* objective function *f*(*x*) over a space of candidates *x*^∗^ = argmax_*x*∈𝒳_ *f*(*x*). Examples of such problems include maximizing the binding affinity of small molecules^10,33^ or proteins^34,35^. Commonly, *f*(*x*) is assumed to be expensive to evaluate and accessible only through evaluation—i.e., the underlying behavior of the objective is unknown.

Bayesian optimization is a sample-efficient model-based framework to solve these costly to evaluate optimization problems^36–38^. At iteration *t* of BO, one has access to observations 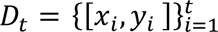 where *y*_*i*_ denotes the (possibly noisy) objective value of the input *x*_*i*_. Typically, a Gaussian process^39^ or approximate Gaussian process^17,40^ is employed as the surrogate model to approximate the objective function using these inputs and values. This surrogate model aids the optimization by employing an acquisition function, which utilizes the surrogate model’s understanding of the objective to strategically propose the next candidates for evaluation. After querying these candidates through the true oracle, the surrogate model is updated with the new observations. This process gradually builds a more comprehensive dataset and refines the surrogate model, thereby improving the quality of the proposed samples in future iterations.

### Bayesian optimization over latent spaces

Due to the discrete and structural nature of peptide sequences, we utilize recent developments in latent space BO that adapt BO from continuous black-box optimization problems to the discrete domain^41,42^. Latent space BO leverages the capabilities of deep generative models, most commonly variational autoencoders (VAEs)^15^ to aid optimization. Concretely, a VAE is composed of two networks: an encoder ℰ (*z*|*x*): 𝒳 → 𝒫(𝒵) mapping from amino acid sequences to a latent space 𝒵, and a decoder 𝒟(*x*|*z*): 𝒵 → 𝒫(𝒳) that probabilistically decodes latent space vectors back into amino acid sequences 𝒳. The VAE is trained in a self-supervised fashion on a large set of unlabeled amino acid sequences using the following standard VAE loss:

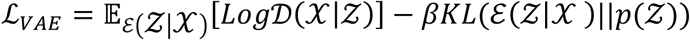

The first term encourages reconstruction accuracy (the amino acid sequences we pass into the encoder being the same as the one we get out of the decoder), and the second KL-divergence term encourages smoothness of the latent space by regularizing the encoder towards the prior distribution *p*(*Z*) ≜ 𝒩(0, 𝑰). In this work, we train a 6-layer transformer encoder and decoder VAE that bottlenecks down to two tokens with 128 latent dimensions each, for a total of 256 latent dimensions. We train with a KL regularization factor of *β* = 10^−4^.

After the VAE is pre-trained on a large number of peptide sequences, for example all sequences in UniRef^43^ within a certain size range, we can define our search over the continuous latent space Z of the VAE instead of the discrete space of amino acid sequences X, we can now formulate our optimization problem as:

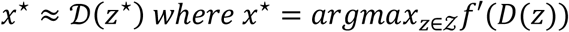

The objective function we are optimizing now takes in a latent vector *z* and decodes it into an amino acid sequence, which is then evaluated on our black box objective function f. Because the true objective function (i.e., MIC) requires expensive lab synthesis to evaluate, we run optimization using our APEX model f’ which estimates the true MIC of the amino acid sequence. After running optimization to obtain sequences that achieve low MICs according to APEX, we then select a batch of the best performing sequences that are then sent to the lab for synthesis and experimental validation. Our optimization algorithm is based on our prior work, the LOL-BO algorithm^10^, which we adapt here for the constrained template derivative optimization setting.

Before beginning optimization, we pre-trained a VAE model on unlabeled amino acid sequences. We train a VAE on 4.5 million amino acid sequences of less than 50 amino acids in length, randomly cropped from the UniRef database^43^ for 118 epochs on a single NVIDIA RTX A6000 GPU to obtain a final test-set reconstruction accuracy of 99.94%. We re-used this same initial pre-trained VAE for all optimization returns, with some additional unlabeled fine-tuning as described below.

We then ran Bayesian optimization in the latent space of the VAE model to find amino acid sequences that minimize *f*′ (*D*(*z*)), where *f*′ was the estimated MIC of the amino acid sequence according to our APEX model. We use a parametric Gaussian process regressor (PPGPR) surrogate with 1024 inducing points to model the data collected during optimization^17^, and select the best candidate latent space point *z* to evaluate next on each iteration of optimization using Thompson sampling to perform acquisition.

Following the LOL-BO algorithm^10^, we also periodically update our VAE model jointly and in and end-to-end fashion with the GP surrogate model to encourage the VAE latent space to re-organize such that peptides with similar scores (MICs) are moved closer together and the space becomes more ideal for modeling and optimization. The models are updated jointly using the following loss:

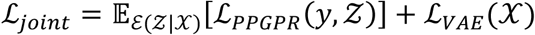

Here, the first term is an expectation over the encoder of the typical^37^ PPGPR loss ℒ_*PPGPR*_(*y*, *Z*), and ℒ_*VAE*_ is the VAE loss as described above.

### Using Trust Regions

In addition to the joint model updates^10^, showed that latent space optimization can be improved by using trust region optimization to limit over-exploration in the very high dimensional latent search space. We therefore use trust region Bayesian optimization^18^. This method works by defining a hyper-rectangular trust region within the search space, and limits the search for new candidates to this local region. The trust region is always defined to be centered on the best point observed so far (the latent space point which achieved the best score *y* = *f*′ (𝒟(𝓏)) we have seen thus far), and the size of the trust region is adapted on each iteration depending on the recent success of the optimizer, with the initial trust region width set to 0.8. If we fail to make progress (to find a new latent point z that achieves a better score than the best we’ve seen so far) for 32 consecutive optimization iterations, the length of the trust region is reduced by a factor of two to further limit overexploitation. Conversely, when we do make progress for 10 consecutive iterations, the length of the trust region doubles to allow more exploration.

### Adding Constraints

A strong desideratum when optimizing each template peptide was to maintain high similarity to the initial template, we choose this similarity constraint because we want to obtain derivatives of peptides that were previously validated, i.e., similarity > 75%. We therefore optimize under an added constraint that all amino acid sequences proposed by the optimization algorithm must be at least 75% similar to the template sequence. Similarity here is measured as (*l* − *d*)/*l*, where *l* is the length of the seed peptide and *d* is the Levenshtein distance between the proposed peptide and the seed.

To add this constraint, we adapted the Scalable Constrained Bayesian Optimization algorithm^44^ to the latent space / generative Bayesian optimization setting. This algorithm works by training a second GP surrogate to model the constraint function 𝒞 (𝒟(𝓏)). In our case, 𝒞 (𝒟(𝓏)) is the similarity of the proposed peptide (𝒟(𝓏) to the template sequence, as measured by fraction of sequence overlap. During optimization, we require that 𝒞 (𝒟(𝓏)) ≥ 75% for all new candidates 𝓏 proposed by the optimizer. On each iteration of optimization, when we use Thompson sampling to select a new candidate point 𝓏 to evaluate next, we exclude all sampled 𝓏𝑠 for which the constraint GP model predicts that the sample will be infeasible (that 𝒞 (𝒟(𝓏𝑠)) < 75%). Then, for each remaining candidate latent vector z, we evaluate the true constraint function 𝒞 (𝒟(𝓏)) and update the constraint GP.

### Fine-tuned VAEs for Template Optimization

Different from the setting considered in the original LOL-BO, we seek to specifically produce optimized derivatives of pre-existing templates because we suspect *a priori* that these de-extinct templates are likely to evade antibiotic resistance. However, VAEs trained on global sets of proteins will devote only small regions of the latent space to producing sequences with high similarity to any particular template: in other words, most sequences produced by the generative model *a priori* would look nothing like the template. To address this, we fine tune the VAE on 20,000 derivatives of the pre-existing template generated via random mutagenesis. This has the effect of significantly increasing the rate at which the VAE produces sequences similar to the template. While the generated sequences will not initially improve antimicrobial activity, this improves gradually over the course of optimization because we update the VAE jointly with the surrogate model.

### Using Multiple Trust Regions to Optimize a Set of Multiple Sequences

Rather than finding a single amino acid sequence that achieves low MIC according to the APEX model, what we want to do is find a set of unique amino acid sequences that all achieve low MIC according to APEX. Having a set of different sequences is important because it increases the chance that some of them will achieve success when they are sent to the wet lab for validation. Thus, rather than searching for a single optimal amino acid sequence, we simultaneously optimize a set of *M* = 20 unique amino acid sequences. To accomplish this, we follow ROBOT^14^ in using 20 different trust regions centered on the 20 unique best-scoring points found so far. Each trust region is responsible for finding one of the 20 final unique sequences in our final set of optimal sequences. On each step of optimization, we select new candidate points from within each of the 20 trust regions. In the *i*th trust region, we discard and resample candidates that are insufficiently diverse from the candidates chosen by trust regions *j* < *i* according to our chosen diversity metric. Specifically, in APEX_GO_, we discard candidates from the i^th^ trust region if any early trust region has already proposed that candidate exactly. We update the size of each trust region according to that individual trust region’s success or failure to propose a new sequence that improves upon the best scoring sequence in that trust region. The 20 trust regions use a shared data history and are re-centered after each iteration so that they are always centered on the best 20 unique sequences observed so far. At the end of the optimization, our final set of optimized sequences are the final centers of the 20 trust regions.

### Optimization goal and setup

During optimization, we aim to produce peptides that achieve low average predicted MIC according to the APEX 1.1 model. We perform separate optimization runs against two objectives: (1) the average predicted MIC against Gram-negative bacteria only (i.e., *A. baumannii* ATCC 19606, *E. coli* ATCC 11775, *E. coli* AIC221, *E. coli* AIC222, *K. pneumoniae* ATCC 13883, *P. aeruginosa* PAO1, *P. aeruginosa* PA14), and (2) against both Gram-negative and Gram-positive bacteria (i.e., *A. baumannii* ATCC 19606, *E. coli* ATCC 11775, *E. coli* AIC221, *E. coli* AIC222, *K. pneumoniae* ATCC 13883, *P. aeruginosa* PAO1, *P. aeruginosa* PA14, *S. aureus* ATCC 12600, methicillin-resistant *S. aureus* ATCC BAA-1556, vancomycin-resistant *E. faecalis* ATCC 700802, and vancomycin-resistant *E. faecium* ATCC 700221). In both cases, the BO minimized the objectives (i.e., lower MICs corresponds to higher antimicrobial activities). In addition to the minimization objective, we used a similarity constraint where each generated peptide should be at most 25% different to given template, where difference is measured as 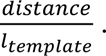 We used a batch size of 50 for optimization, with 20 trust regions and *τ* = 1, meaning that we aim to produce 20 different peptides for the same template in a single optimization run. We update our surrogate model and VAE jointly every 10 steps of optimization and initialize the optimization with 10,000 peptides randomly mutated from the template peptide.

### Peptide Synthesis

All peptides for the experiments were obtained from AAPPTec and synthesized using solid-phase peptide synthesis with the Fmoc strategy.

### Minimal inhibitory concentration assays

Broth microdilution assays were conducted to establish the minimum inhibitory concentration (MIC) for each peptide^9,45,46^. Peptides were added to untreated polystyrene 96-well microtiter plates and serially diluted two-fold in sterile water, ranging from 0 to 64 μmol L^-1^. A bacterial inoculum at a concentration of 10⁶ CFU mL^-1^ in LB medium was then mixed in a 1:1 ratio with the peptide solution. The MIC was determined as the lowest peptide concentration that completely inhibited bacterial growth after 24 h of incubation at 37 °C. Each assay was performed in three independent replicates.

### Circular dichroism experiments

The circular dichroism experiments were conducted using a J1500 circular dichroism spectropolarimeter (Jasco) in the Biological Chemistry Resource Center (BCRC) at the University of Pennsylvania. Experiments were performed at 25 °C, the spectra graphed are an average of three accumulations obtained with a quartz cuvette with an optical path length of 1.0 mm, ranging from 260 to 190 nm at a rate of 50 nm min^-1^ and a bandwidth of 0.5 nm. The concentration of all EPs tested was 50 μmol L^-1^, and the measurements were performed in water, a mixture of trifluoroethanol (TFE) and water in a 3:2 ratio, and sodium dodecyl sulfate (SDS) in water at 10 mmol L^-1^, with respective baselines recorded prior to measurement. A Fourier transform filter was applied to minimize background effects. Secondary structure fraction values were calculated using the single spectra analysis tool on the server BeStSel^28^. Ternary plots were created in https://www.ternaryplot.com/ and subsequently edited.

### Outer membrane permeabilization assays

N-phenyl-1-napthylamine (NPN) uptake assay was used to evaluate the ability of the peptides to permeabilize the bacterial outer membrane. Inocula of *A. baumannii* ATCC 19606 were grown to an OD at 600 nm of 0.4 mL^-1^, centrifuged (10,000 rpm at 4 °C for 10 min), washed and resuspended in 5 mmol L^-1^ HEPES buffer (pH 7.4) containing 5 mmol L^-1^ glucose. The bacterial solution was added to a white 96-well plate (100 μL per well) together with 4 μL of NPN at 0.5 mmol L^-1^. Consequently, peptides diluted in water were added to each well, and the fluorescence was measured at λ_ex_ = 350 nm and λ_em_ = 420 nm over time for 45 min. The relative fluorescence was calculated using the untreated control (buffer + bacteria + fluorescent dye) and polymyxin B (positive control) as baselines and the following equation was applied to reflect % of difference between the baselines and the sample:

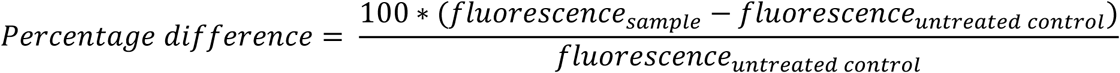

### Cytoplasmic membrane depolarization assays

The cytoplasmic membrane depolarization assay was performed using the membrane potential-sensitive dye 3,3’-dipropylthiadicarbocyanine iodide (DiSC_3_-5). *A. baumannii* ATCC 19606 in the mid-logarithmic phase were washed and resuspended at 0.05 OD mL^-1^ (optical value at 600 nm) in HEPES buffer (pH 7.2) containing 20 mmol L^-1^ glucose and 0.1 mol L^-1^ KCl. DiSC3-5 at 20 μmol L^-1^ was added to the bacterial suspension (100 μL per well) for 15 min to stabilize the fluorescence which indicates the incorporation of the dye into the bacterial membrane, and then the peptides were mixed 1:1 with the bacteria to a final concentration corresponding to their MIC values. Membrane depolarization was then followed by reading changes in the fluorescence (λ_ex_ = 622 nm, λ_em_ = 670 nm) over time for 60 min. The relative fluorescence was calculated using the untreated control (buffer + bacteria + fluorescent dye) and polymyxin B (positive control) as baselines and the following equation was applied to reflect % of difference between the baselines and the sample:

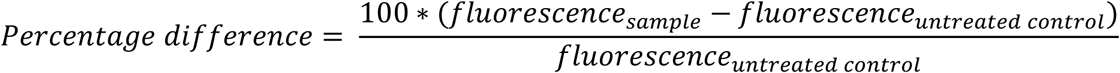

### Cytotoxicity assays

One day before the experiment, an aliquot of 100 μL of the cells at 50,000 cells per mL was seeded into each well of the cell-treated 96-well plates used in the experiment. The attached HEK293T cells were then exposed to increasing concentrations of the peptides (8–128 μmol L^-1^) for 24 h. After the incubation period, we performed the 3-(4,5-dimethylthiazol-2-yl)-2,5-diphenyltetrazolium bromide tetrazolium reduction assay (MTT assay)^9^. The MTT reagent was dissolved at 0.5 mg mL^-1^ in medium without phenol red and was used to replace cell culture supernatants containing the peptides (100 μL per well), and the samples were incubated for 4 h at 37 °C in a humidified atmosphere containing 5% CO_2_ yielding the insoluble formazan salt. The resulting salts were then resuspended in hydrochloric acid (0.04 mol L^-1^) in anhydrous isopropanol and quantified by spectrophotometric measurements of absorbance at 570 nm. All assays were done as three biological replicates.

### Skin abscess infection mouse model

The back of six-week-old female CD-1 mice under anesthesia were shaved and injured with a superficial linear skin abrasion made with a needle. An aliquot of *A. baumannii* ATCC 19606 (5×10^5^ CFU mL^-1^; 20 μL) previously grown in LB medium until 0.5 OD mL^-1^ (optical value at 600 nm) and then washed twice with sterile PBS (pH 7.4, 13,000 rpm for 3 min) was added to the scratched area. Peptides diluted in sterile water at their MIC value were administered to the wounded area 1 h post-infection. Two- and four-days post-infection, animals were euthanized, and the scarified skin was excised, homogenized using a bead beater (25 Hz for 20 min), 10-fold serially diluted, and plated on McConkey agar plates for CFU quantification. The experiments were performed using four mice per group. The skin abscess infection mouse model was revised and approved by the University Laboratory Animal Resources (ULAR) from the University of Pennsylvania (Protocol 806763).

### Deep thigh infection mouse model

Experiments were performed using six-week-old female CD-1 mice, which were rendered neutropenic by intraperitoneal application of two doses of cyclophosphamide (150 mg Kg^-1^ and 100 mg Kg^-1^) 3 and 1 days before the infection. At day 4 of the experiment, the mice were infected in their right thigh through a 100 μL intramuscular injection of *A. baumannii* ATCC19606 (in PBS at a concentration of 9×10^5^ CFU mL^-1^). The bacterial cells were grown in LB broth, washed twice with PBS solution, and diluted at the desired concentration prior to infecting the mice. The peptides were administered intraperitoneally two hours after the infection. Four-days post-infection mice were euthanized, and the tissue from the right thigh was excised, homogenized using a bead beater (25 Hz for 20 min), 10-fold serially diluted, and plated on McConkey agar plates for bacterial colonies counting. The experiments were performed using four mice per group. The deep thigh infection mouse model was revised and approved by the University Laboratory Animal Resources (ULAR) from the University of Pennsylvania (Protocol 807055).

## Acknowledgments

Cesar de la Fuente-Nunez holds a Presidential Professorship at the University of Pennsylvania and acknowledges funding from the Procter & Gamble Company, United Therapeutics, a BBRF Young Investigator Grant, the Nemirovsky Prize, Penn Health-Tech Accelerator Award, and the Dean’s Innovation Fund from the Perelman School of Medicine at the University of Pennsylvania. Research reported in this publication was supported by the Langer Prize (AIChE Foundation), the National Institute of General Medical Sciences of the National Institutes of Health under award number R35GM138201, and the Defense Threat Reduction Agency (DTRA; HDTRA11810041, HDTRA1-21-1-0014, and HDTRA1-23-1-0001). We thank Dr. Mark Goulian for kindly donating the following strains: *Escherichia coli* AIC221 [*Escherichia coli* MG1655 phnE_2::FRT (control strain for AIC 222)] and *Escherichia coli* AIC222 [*Escherichia coli* MG1655 pmrA53 phnE_2::FRT (polymyxin resistant)]. We thank de la Fuente Lab members for insightful discussions. Figures created with BioRender.com are attributed as such. Molecules were rendered using the PyMOL Molecular Graphics System, Version 3.0 Schrödinger, LLC.

## Author contributions

Conceptualization: MDTT, YZ, FW, NM, JG, CFN

Methodology: MDTT, YZ, FW, NM

Computational investigation: YZ, FW, NM

Experimental investigation: MDTT

Visualization: MDTT, YZ, FW, NM

Funding acquisition: JG, CFN

Supervision: JG, CFN

Software: YZ, FW, NM, JG

Formal analysis: MDTT, YZ, FW, NM, JG

Writing – original draft: MDTT, YZ, FW, NM, JG, CFN

Writing – review & editing final version of the paper: MDTT, YZ, FW, NM, JG, CFN

## Data and materials availability

The APEX_GO_ model is available on Github (https://github.com/Yimeng-Zeng/APEXGo). All data pertaining to the experimental validation of generated peptides are available in the Supplementary Data and Mendeley Data (https://data.mendeley.com/datasets/kk53h8pt5v/1).

## Supplementary Information

**Supplementary Table 1.**
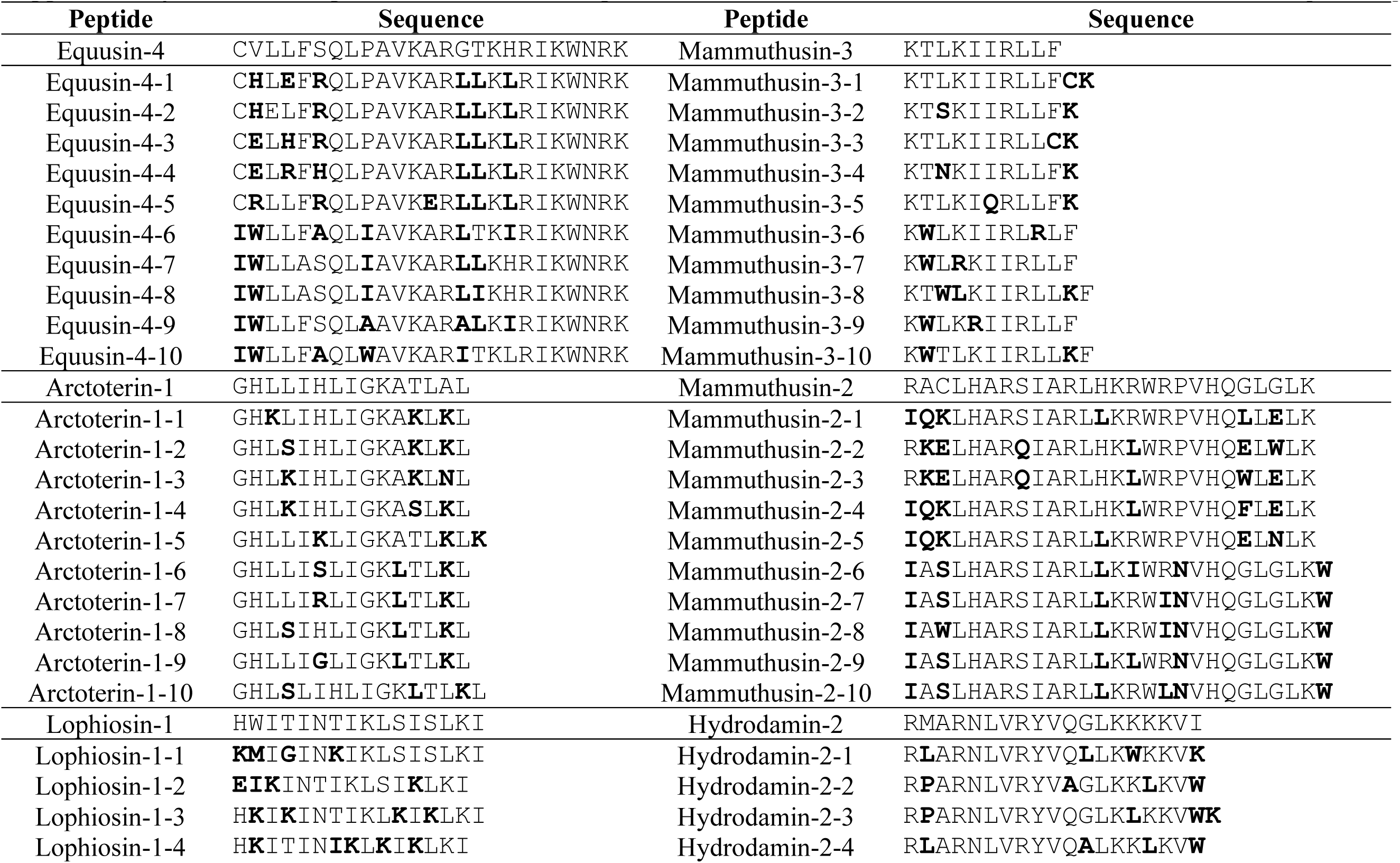
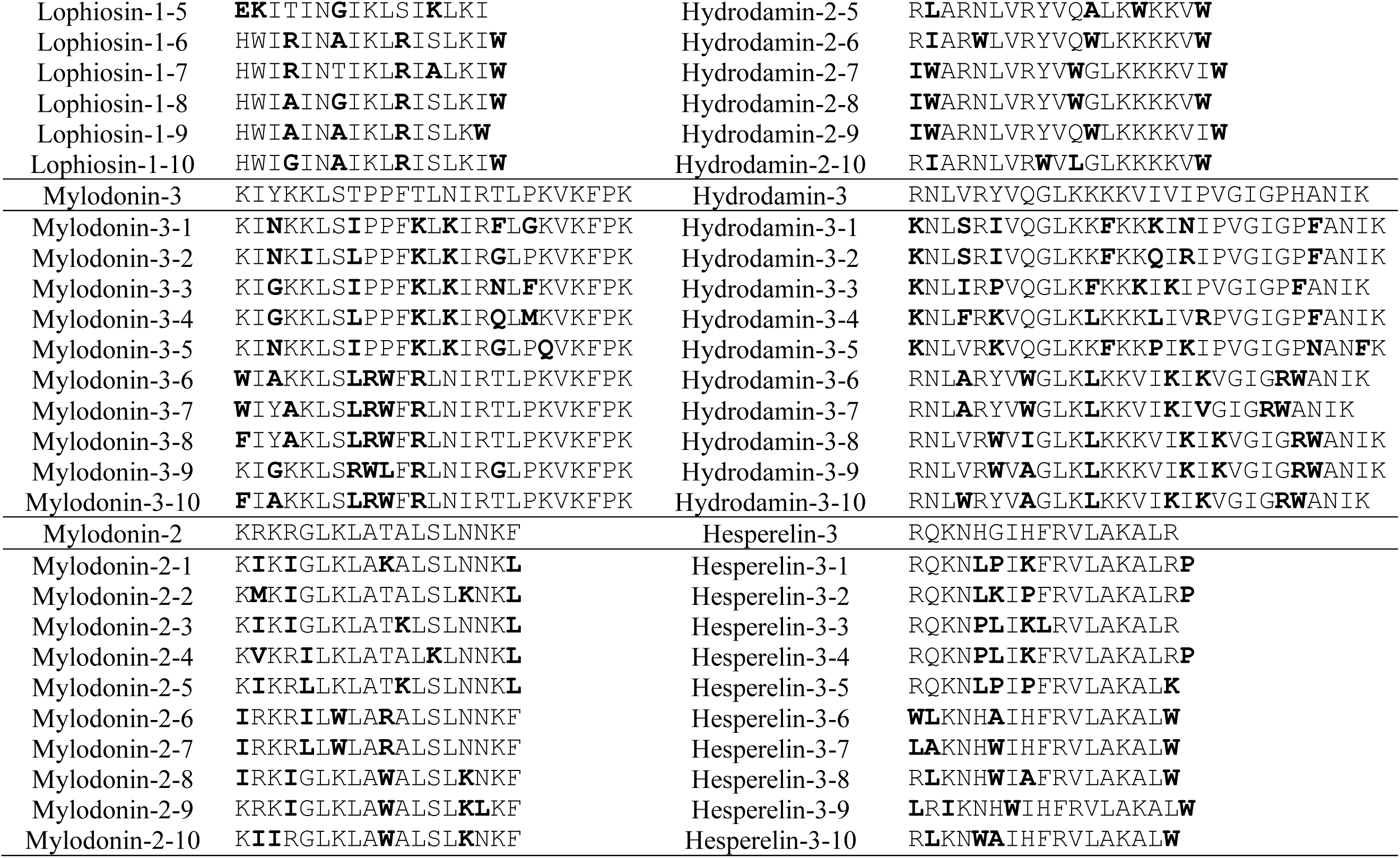
Template and derivatives sequences. In bold amino acid residues that are different from the original sequence.

**Supplementary Figure 1.**
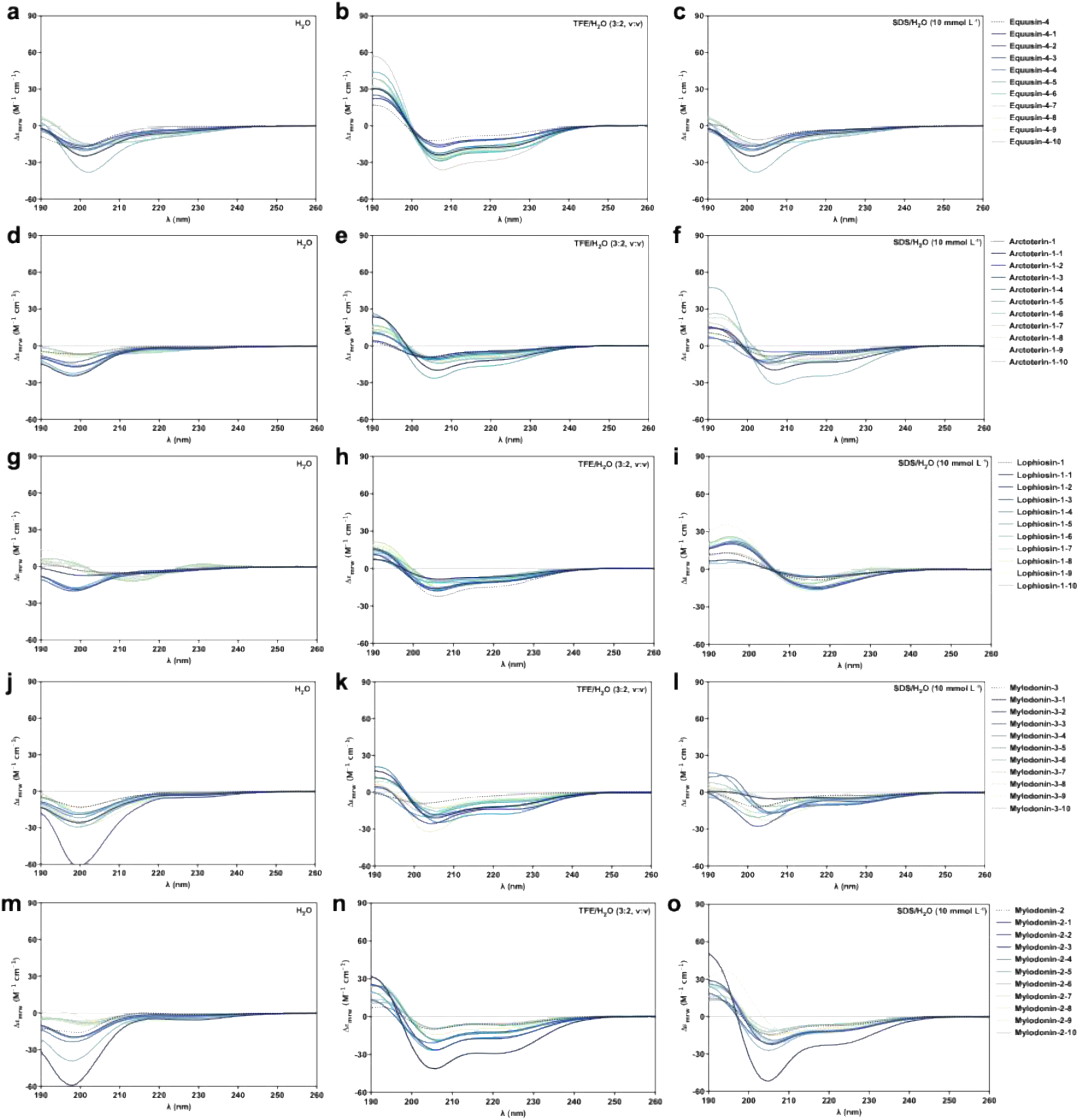
Circular dichroism spectra of templates equusin-4, arctoterin-1, lophiosin-1, mylodonin-3, and mylodonin-2 and their respective APEX_GO_-optimized derivatives. Circular dichroism experiments were conducted in a J-1500 Jasco circular dichroism spectrophotometer. The spectra were recorded in three different media: water, 60% trifluoroethanol in water, and Sodium dodecyl sulfate (SDS) in water (10 mmol L^-1^), after three accumulations at 25 °C, using a 1mm path length quartz cell, between 260 and 190 nm at 50 nm min^-1^, with a bandwidth of 0.5 nm. The concentration of all peptides tested was 50 μmol L^-1^.

**Supplementary Figure 2.**
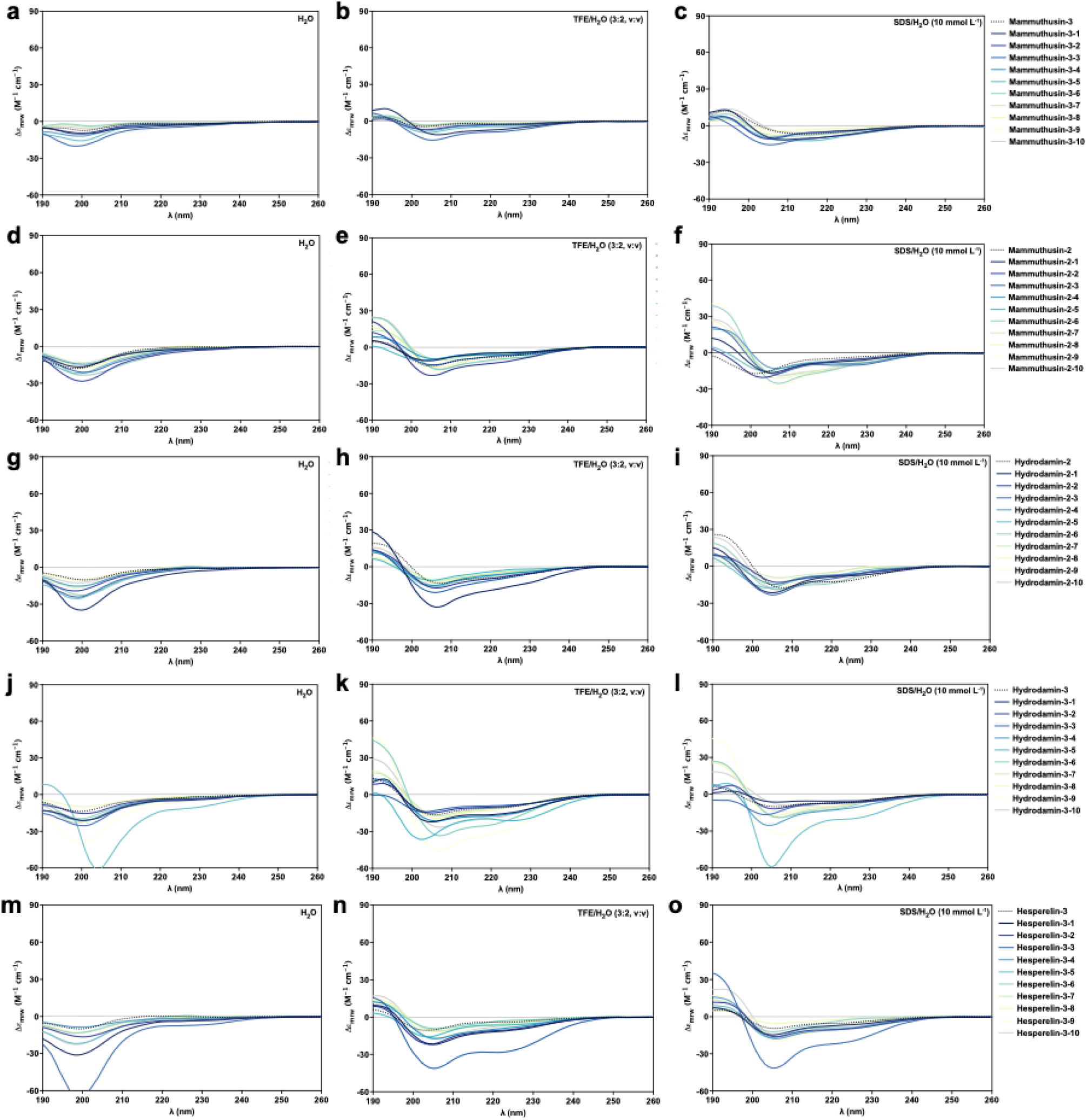
Circular dichroism spectra of templates mammuthusin-3, mammuthusin-2, hydrodamin-2, hydrodamin-3, and hesperelin-3 and their respective APEX_GO_-optimized derivatives. Circular dichroism experiments were conducted in a J-1500 Jasco circular dichroism spectrophotometer. The spectra were recorded in three different media: water, 60% trifluoroethanol in water, and Sodium dodecyl sulfate (SDS) in water (10 mmol L^-1^), after three accumulations at 25 °C, using a 1mm path length quartz cell, between 260 and 190 nm at 50 nm min^-1^, with a bandwidth of 0.5 nm. The concentration of all peptides tested was 50 μmol L^-1^.

**Supplementary Figure 3.**
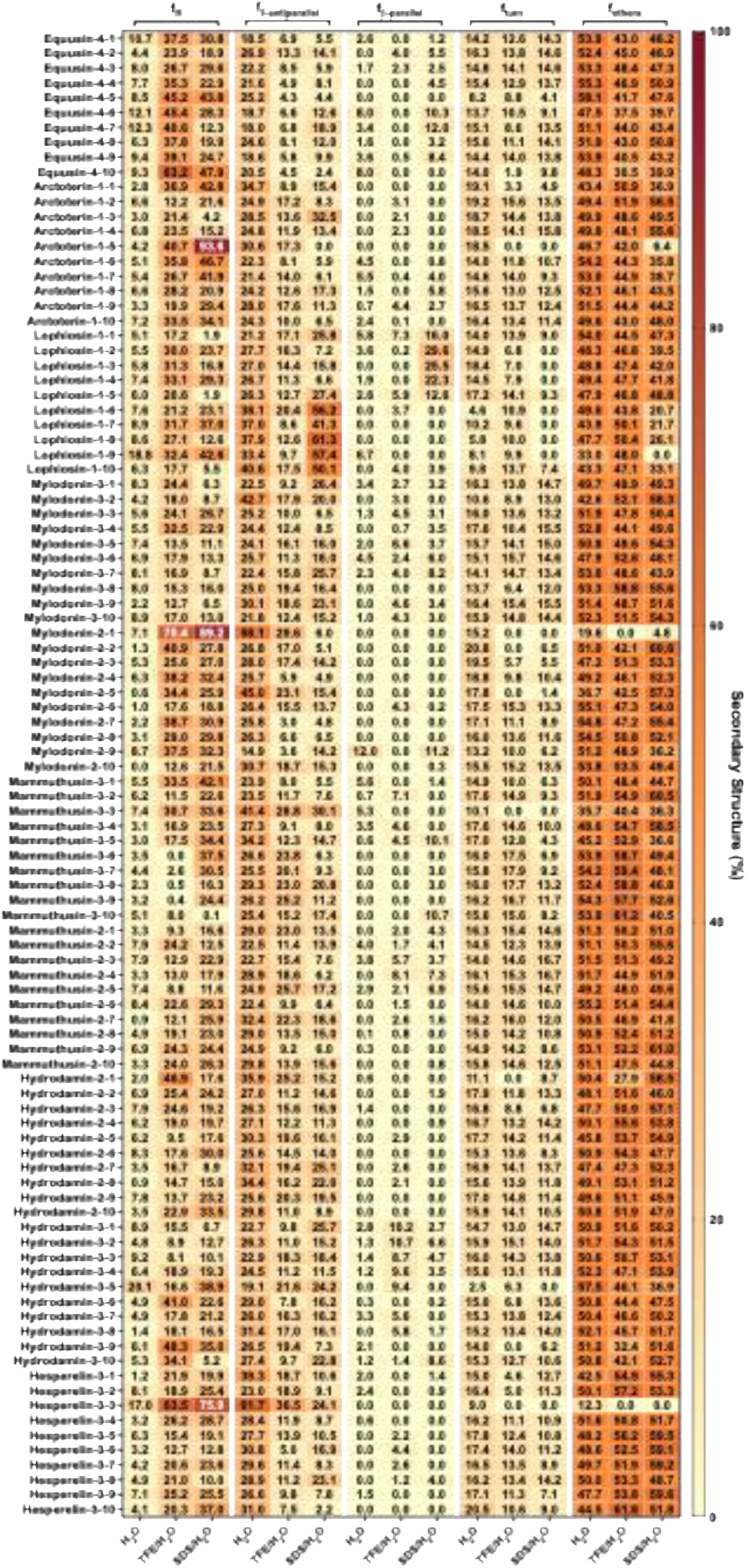
Secondary structure fractions of APEX_GO_-optimized peptides. Heatmap with the percentage of secondary structure found for each peptide in three different solvents: water, 60% trifluoroethanol (TFE) in water, and SDS (10 mmol L^-1^) in water. Secondary structure fraction was calculated using the BeStSel server^28^.

**Supplementary Figure 4.**
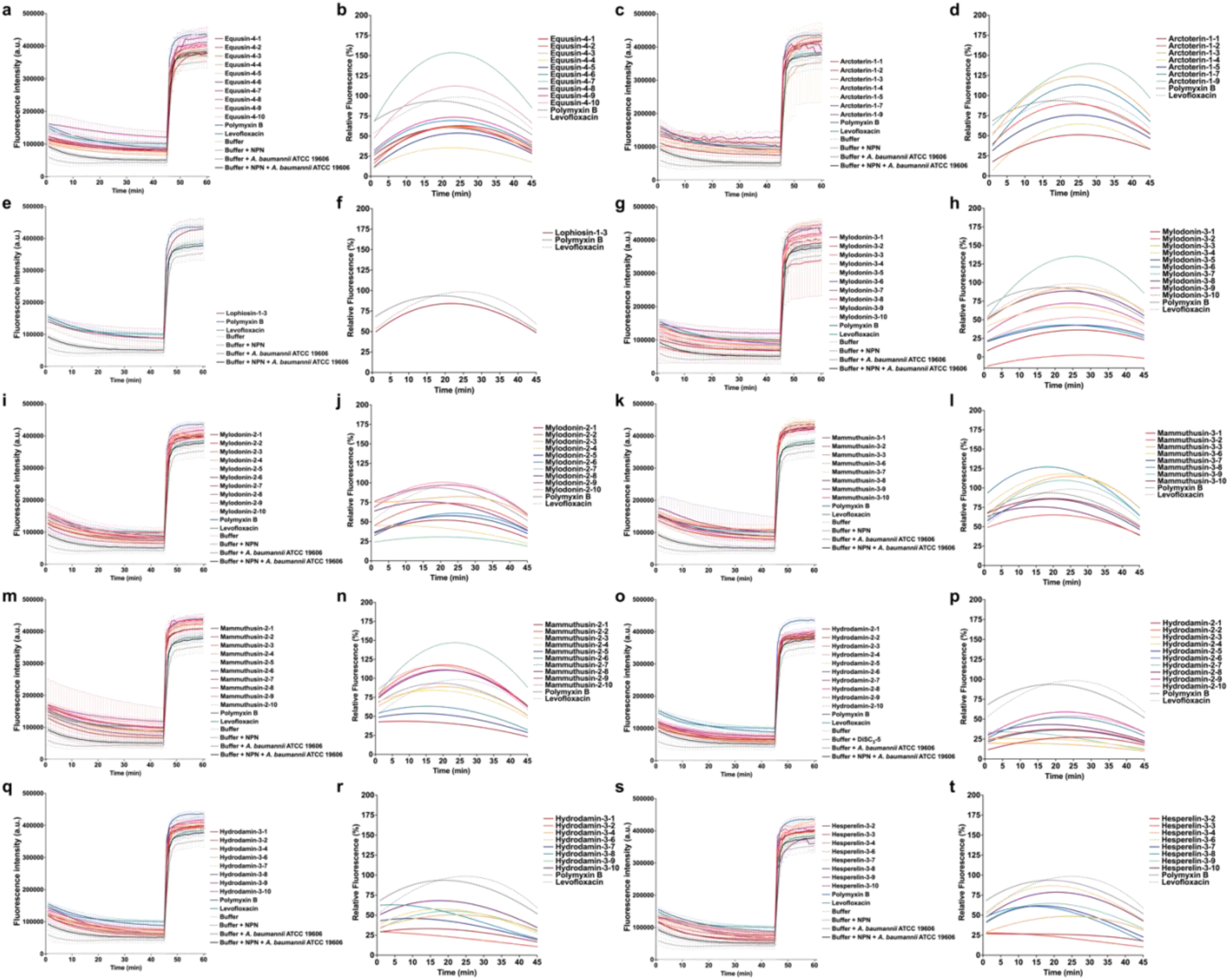
Outer membrane permeabilization of *A. baumannii* ATCC 19606 induced by APEX_GO_-optimized peptides. Outer membrane permeabilization was assessed using the probe 1-(N-phenylamino)naphthalene (NPN), showing the permeabilization effects of peptides derived from each of the following 10 parent de-extinct EPs that showed antimicrobial activity against *A. baumannii* ATCC 19606: **(a)** equusin-4, **(c)** arctoterin-1, **(e)** lophiosin-1, **(g)** mylodonin-3, **(i)** mylodonin-2, **(k)** mammuthusin-3, **(m)** mammuthusin-2, **(o)** hydrodamin-2, **(q)** hydrodamin-3, and **(s)** hesperelin-3. The relative fluorescence values of the probe’s fluorescence obtained for each of the experiments and represented by a non-linear fitting compared to the baseline of the untreated control (buffer + bacteria + fluorescent dye) and benchmarked against the antibiotics polymyxin B and levofloxacin are represented for: **(b)** equusin-4, **(d)** arctoterin-1, **(f)** lophiosin-1, **(h)** mylodonin-3, **(j)** mylodonin-2, **(l)** mammuthusin-3, **(n)** mammuthusin-2, **(p)** hydrodamin-2, **(r)** hydrodamin-3, and **(t)** hesperelin-3.

**Supplementary Figure 5.**
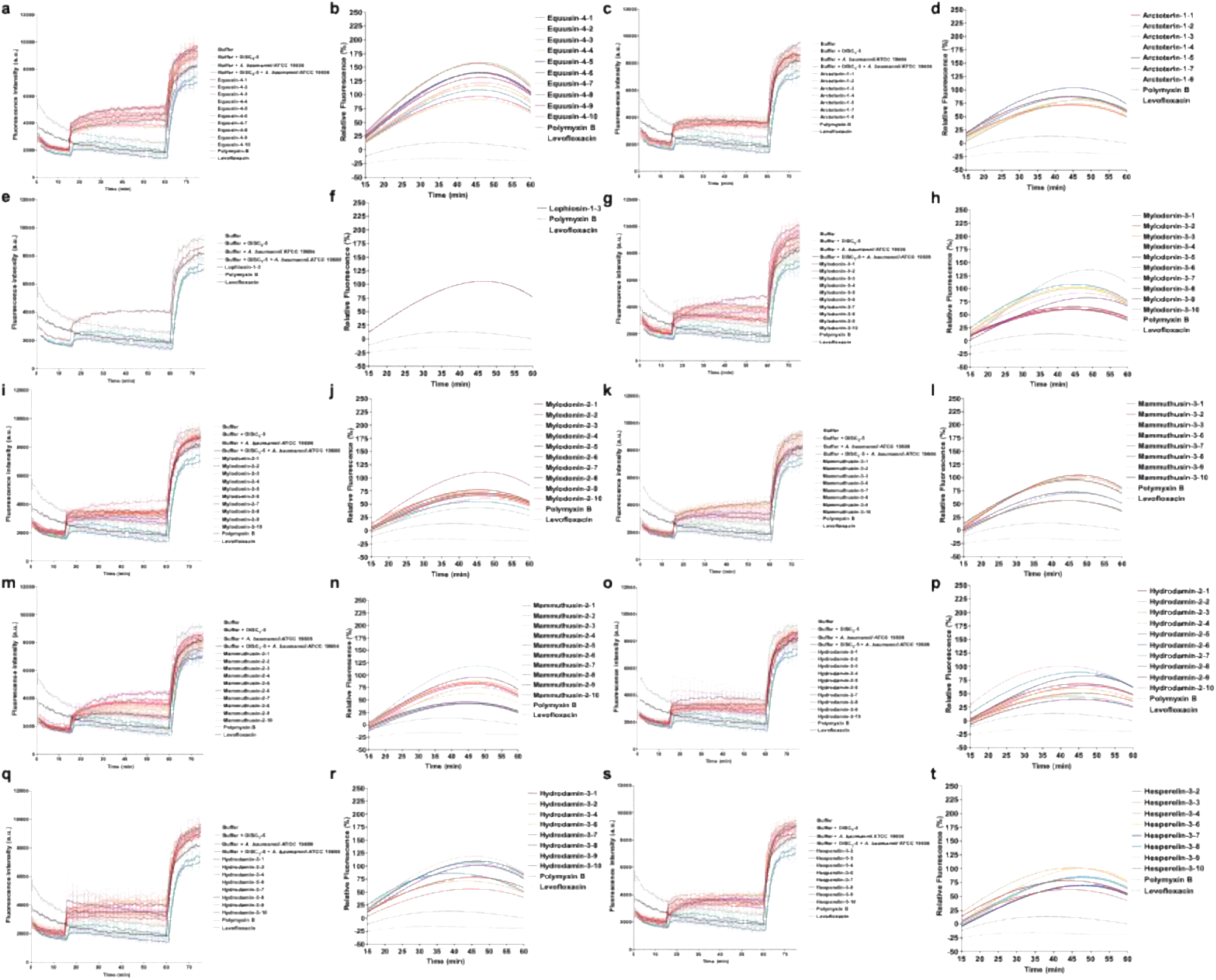
Outer membrane permeabilization and cytoplasmic membrane depolarization of *A. baumannii* ATCC 19606 induced by APEX_GO_-optimized peptides. Membrane depolarization assays were performed using the hydrophobic probe 3,3′-dipropylthiadicarbocyanine iodide [DiSC_3_-(5)] showing the depolarization effects of APEX_GO_-optimized peptides derived from each of the following 10 parent de-extinct peptides that showed antimicrobial activity against *A. baumannii* ATCC 19606: **(a)** equssin-4, **(c)** arctoterin-1, **(e)** lophiosin-1, **(g)** mylodonin-3, **(i)** mylodonin-2, **(k)** mammuthusin-3, **(m)** mammuthusin-2, **(o)** hydrodamin-2, **(q)** hydrodamin-3, and **(s)** hesperelin-3. The relative fluorescence values of the probe’s fluorescence obtained for each of the experiments and represented by a non-linear fitting compared to the baseline of the untreated control (buffer + bacteria + fluorescent dye) and benchmarked against the antibiotics polymyxin B and levofloxacin are represented for: **(b)** equssin-4, **(d)** arctoterin-1, **(f)** lophiosin-1, **(h)** mylodonin-3, **(j)** mylodonin-2, **(l)** mammuthusin-3, **(n)** mammuthusin-2, **(p)** hydrodamin-2, **(r)** hydrodamin-3, and **(t)** hesperelin-3.

**Supplementary Figure 6.**
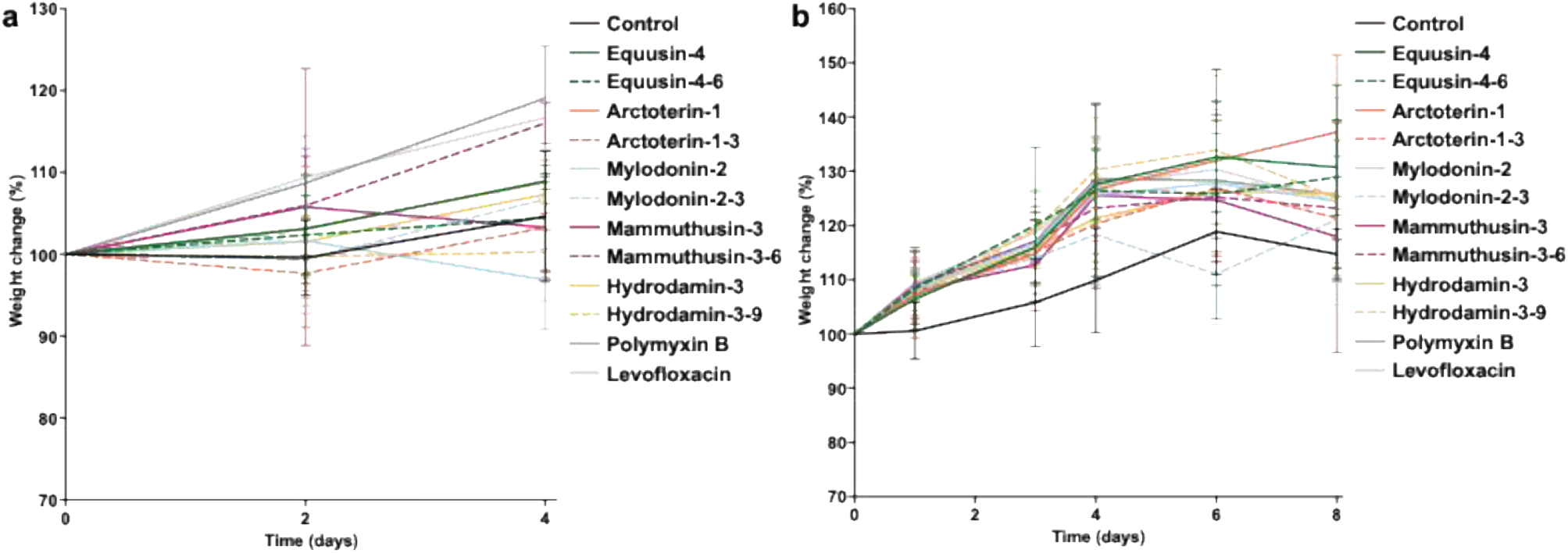
Weight change monitoring in both skin abscess and deep thigh infection mouse models infected with *A. baumannii*. Mouse weight was monitored throughout the duration of the **(a)** skin abscess model (4 days total) and the **(b)** deep thigh infection model (8 days total) to assess potential toxic effects of both the bacterial load and the peptides.

